# Metabolic dependency of chorismate in *Plasmodium falciparum*

**DOI:** 10.1101/698951

**Authors:** Ana Lisa Valenciano, Maria L. Fernández-Murga, Emilio F. Merino, Nicole R. Holderman, Grant J. Butschek, Karl J. Shaffer, Peter C. Tyler, Maria Belen Cassera

**Author notes:** To whom correspondence should be addressed: Maria Belen Cassera: Department of Biochemistry & Molecular Biology, University of Georgia, Athens GA 30602.

## Abstract

The shikimate pathway, a metabolic pathway absent in humans, is responsible for the production of chorismate, a branch point metabolite. In the malaria parasite, chorismate is postulated to be a direct precursor in the synthesis of *p*-aminobenzoic acid (folate biosynthesis), *p*-hydroxybenzoic acid (ubiquinone biosynthesis), menaquinone, and aromatic amino acids. While the potential value of the shikimate pathway as a drug target is debatable, the metabolic dependency of chorismate in *P. falciparum* remains unclear. Current evidence suggests that the main role of chorismate is folate biosynthesis despite ubiquinone biosynthesis being active and essential in the malaria parasite. Our goal in the present work was to expand our knowledge of the ubiquinone head group biosynthesis and its potential metabolic dependency on chorismate in *P. falciparum.* These data led us to further characterize the mechanism of action of MMV688345, a compound from the open-access “Pathogen Box” collection from Medicine for Malaria Venture. We systematically assessed the development of both asexual and sexual stages of *P. falciparum* in a defined medium in the absence of an exogenous supply of chorismate end-products and present biochemical evidence suggesting that the benzoquinone ring of ubiquinones in this parasite may be synthesized through a yet unidentified route.

---

Human malaria accounts for more than 400,000 deaths every year, with *Plasmodium falciparum* being the deadliest of the species that infects humans ^1^. All *Plasmodium* species have a complex life cycle that occurs between the human host and the *Anopheles* mosquito vector. Clinical symptoms are caused by the asexual intraerythrocytic cycle of the parasite that lasts around 48 h in *P. falciparum*. During the asexual cycle, the parasite progresses through four morphologically different stages: ring, trophozoite, and schizont stages, ending with the release of merozoites that will invade new erythrocytes. A small percentage (0.1-0.2) of parasites commit to gametocytogenesis (sexual development) during the asexual cycle and mature female and male gametocytes are transmitted to a female mosquito when it feeds on an infected human. Gametocytes are asymptomatic, non-replicating forms that can persist for weeks in circulating blood. In contrast to other species of *Plasmodium, P. falciparum* gametocytes develop through five (I to V) morphologically distinct stages, taking 10 to 12 days to fully mature into stage V gametocytes.

Artemisinin Combination Therapies (ATCs) are the frontline treatment for malaria, however, resistance to artemisinin has been confirmed and is increasing in prevalence ^2^. Thus, there is an urgent need to identify novel drugs against new therapeutic targets. Metabolic pathways have been a source of druggable targets across many human diseases, especially for infectious diseases when pathways or their enzymes are sufficiently different or absent in the human host as is the case for the shikimate pathway ^3^.

The shikimate pathway consists of seven enzymatic steps for production of chorismate, a branch point metabolite that is then used for the synthesis of a diverse range of end products, depending on the organism (Fig. 1). In the malaria parasite, chorismate is postulated to be a direct precursor in the synthesis of *p*-aminobenzoic acid (pABA, an intermediate of folate biosynthesis), *p*-hydroxybenzoic acid (pHBA, a precursor of ubiquinone biosynthesis), menaquinone, and aromatic amino acids (tryptophan, phenylalanine and tyrosine) ^4-6^. However, few studies have presented experimental evidence regarding the metabolic dependency of the malaria parasite on the shikimate pathway. Roberts and colleagues demonstrated that *P. falciparum in vitro* growth inhibition by glyphosate, a known 5-enolpyruvylshikimate 3-phosphate synthase (EPSPS) inhibitor often used as herbicide, could be reversed by supplementing the media with pABA. This suggested the presence of an EPSPS in the malaria parasite and provided evidence of its role in folic acid biosynthesis ^7^. In addition, the effect of fluorinated analogs of shikimate on growth of *P. falciparum* in media deficient in aromatic metabolites has been explored by McConkey where the inhibitory growth effect was reversed by exogenous supplementation of pHBA, pABA, and aromatic amino acids (tryptophan, tyrosine and phenylalanine) ^3^. However, the contribution of the shikimate pathway to the pool of tryptophan, tyrosine and phenylalanine is difficult to assess since these amino acids are also obtained from hemoglobin digestion ^8^.

**Figure 1.**
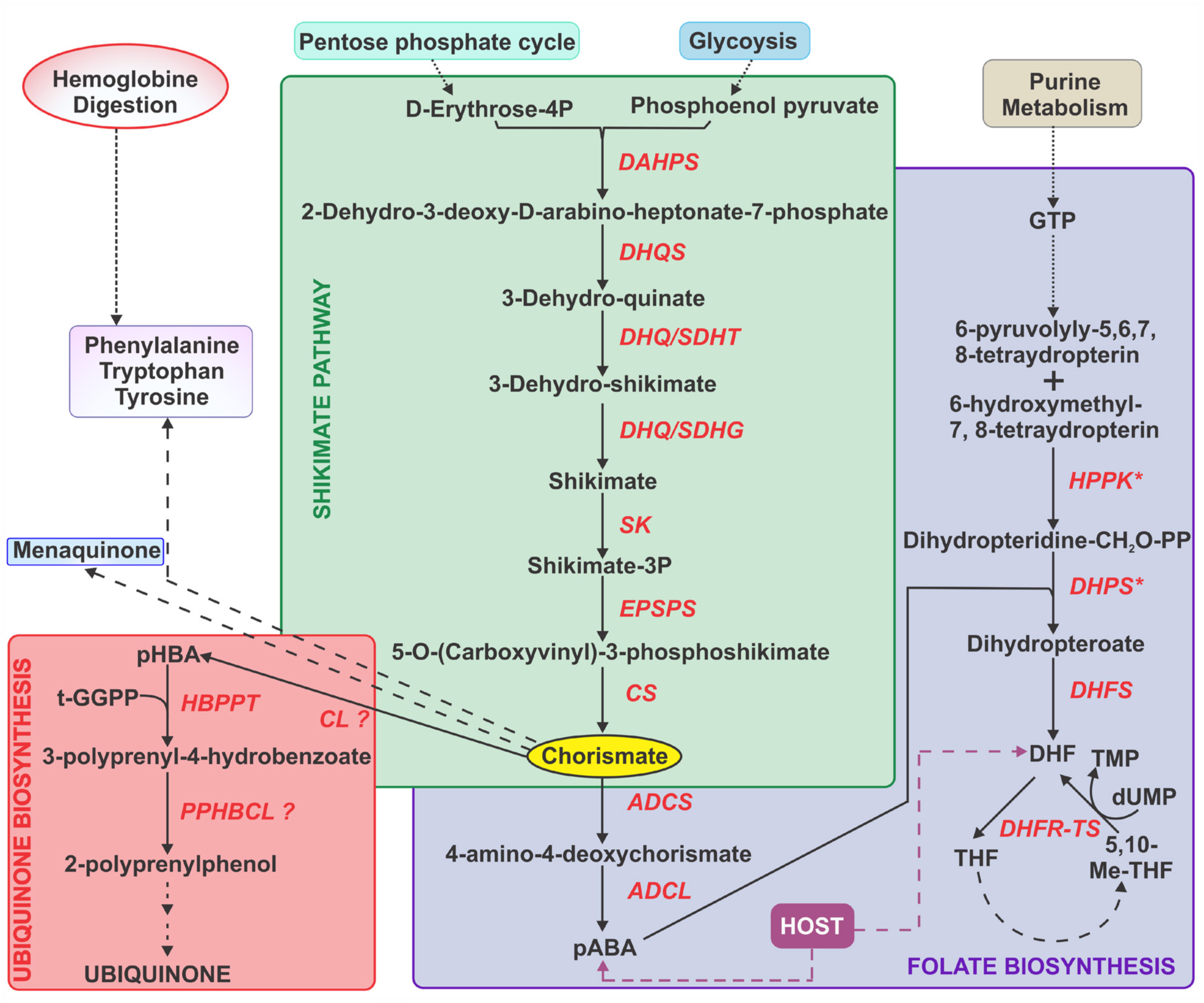
Postulated chorismate dependency in *P. falciparum.* The shikimate pathway consists of seven enzymatic steps for production of chorismate, a branch point metabolite: DAHPS, 3-deoxy-7-phosphoheptulonate synthase; DHQS, 3-dehydroquinate synthase; DHQ/SDHT, 3-dehydroquinate dehydratase; DHQ/SDHG, shikimate 5-dehydrogenase; SK, shikimate kinase; EPSPS, 5-enolpyruvylshikimate 3-phosphate synthase; CS, chorismate synthase. Ubiquinone biosynthesis: CL, chorismate lyase; HBPPT, 4-hydroxybenzoate polyprenyltransferase; PPHBCL, 3-polyprenyl-4-hydroxybenzoate carboxy-lyase. Folate biosynthesis: ADCS, aminodeoxy chorismate synthase; ADCL, aminodeoxy chorismate lyase; HPPK-DHPS, bifunctional 6-hydroxymethyl-7,8-dihydropterin pyrophosphokinase/dihydropteroate synthase; DHFS, dihydrofolate synthase; DHFR-TS, bifunctional dihydrofolate reductase/thymidylate synthase. Both folate, that is converted to dihydrofolate, and pABA can be salvaged from the host.

Though seven enzymes are postulated to be involved in the chorismate biosynthesis, only chorismate synthase has been clearly identified and studied in *P. falciparum* ^5,9^. In addition, bioinformatic analysis suggested the presence of a potential candidate for shikimate 5-dehydrogenase (PF3D7-1444700) in *P. falciparum* ^10^. Remarkably, a recent study revealed that chorismate synthase (CS) is not essential for *P. berghei* to survive throughout the parasite’s life cycle ^11^. Additionally, two recent functional profiling studies of the *Plasmodium* genome also indicated that CS may be dispensable in asexual stages of *P. berghei* in mice and *in vitro P. falciparum* ^12,13^. These results may be due to the presence of pABA in the mice’s diet as well as in the culture media used in the *in vitro* assays for *P. falciparum*. Low levels of pABA have been reported as unfavorable for *Plasmodium* growth *in vitro* and *in vivo* in different malaria models including *P. falciparum-*infected *Aotus* monkeys (reviewed in ^14^). Thus, this metabolite has been considered a necessary exogenous nutrient to support parasite growth.

While the potential value of the shikimate pathway as a drug target is still debatable and several genes have yet to be identified, the metabolic dependency of chorismate in *P. falciparum* remains unknown. The current available research described above suggests that the main role of chorismate is folate biosynthesis despite active ubiquinone biosynthesis in the malaria parasite ^15,16^. Ubiquinone biosynthesis is postulated to be essential in *Plasmodium* asexual stages based on recent functional profiling studies of *Plasmodium* genome ^12,13^. However, similarly to the shikimate pathway, no genes can be found for chorismate lyase (CL, EC 4.1.3.40), the first enzyme of ubiquinone biosynthesis, which converts chorismate into pHBA or the enzyme 3-polyprenyl-4-hydroxybenzoate carboxy-lyase (PPHBCL) responsible for *C*1-decarboxylation of the head group (Fig. 1).

One of the challenges of studying metabolic pathways in *P. falciparum* is that while its genome being sequenced in 2002 ^17^, around 45% of the genes remain annotated as hypothetical proteins without a known function. The “Malaria Parasite Metabolic Pathways” (MPMP) ^18^ and “Library of Apicomplexan Metabolic Pathways” (LAMP) ^19^ databases are the only manually curated databases available for Apicomplexa using both genomic and biochemical/physiological evidence. Therefore, our goal in the present work was to expand our knowledge of the ubiquinone head group biosynthesis and its potential metabolic reliance on chorismate in *P. falciparum*. Our investigations, led us to further characterize a compound previously reported as antifolate present in the open-access “Pathogen Box” collection (MMV688345) from Medicine for Malaria Venture (MMV) ^20,21^. To our knowledge, this is the first systematic study assessing the metabolic dependency of chorismate in *P. falciparum*.

## Results

### Development of P. falciparum asexual and gametocyte stages in the absence of chorismate end-products found in standard growth medium

Our first step was to systematically assess the development of both asexual and sexual stages of *P. falciparum* in a defined medium in the absence of an exogenous supply of chorismate end-products. It has been previously reported that asexual stages of *P. falciparum* can grow in pABA- and folate-deficient medium ^22-25^, which forces parasites to rely on their own biosynthesis using the shikimate pathway. Therefore, in order to assess the metabolic dependency of chorismate for ubiquinone biosynthesis, we sought to standardize and characterize *P. falciparum* asexual growth and sexual development in a defined media, here referred to as RPMI Minimal Medium (MM), lacking aromatic metabolites (pABA, folic acid (FA), tryptophan, phenylalanine and tyrosine) that are found in the standard RPMI medium (CM) (Table S1). *P. falciparum* was adapted to grow in MM for at least two weeks before tests began and were maintained for at least 3 months. As shown in Fig. 2A, parasites grown in MM (asexual cycle) did not show any morphological differences compared to those grown in CM. However, a lower percentage of ring stages in the MM condition was consistently detected after three independent analyses (Fig. 2B). Consistent with previous reports (reviewed in ^14^), supplementation of MM with FA at 2.2 μM or pABA at 7.3 μM resulted in a similar percentage of rings compared to CM. Interestingly, a similar result was obtained with chorismate indicating that exogenous chorismate supplementation may increase folate biosynthesis which in turn supports better *in vitro* growth of *P. falciparum*. Altogether, these results suggest that folate or its derivatives may play a role during egress/invasion of merozoites and ring development. Further investigation is required as pABA is the only known metabolite present in adult human plasma at variable concentrations (0.15 - 6.56 μM) ^26^ and increasing folate supplementation in malaria endemic areas ^27^ has the potential to result in an increased risk of infection and failure of antifolate treatments. It is worth mentioning that all metabolites except pABA are present in the CM at supraphysiological concentrations ^28^.

**Figure 2.**
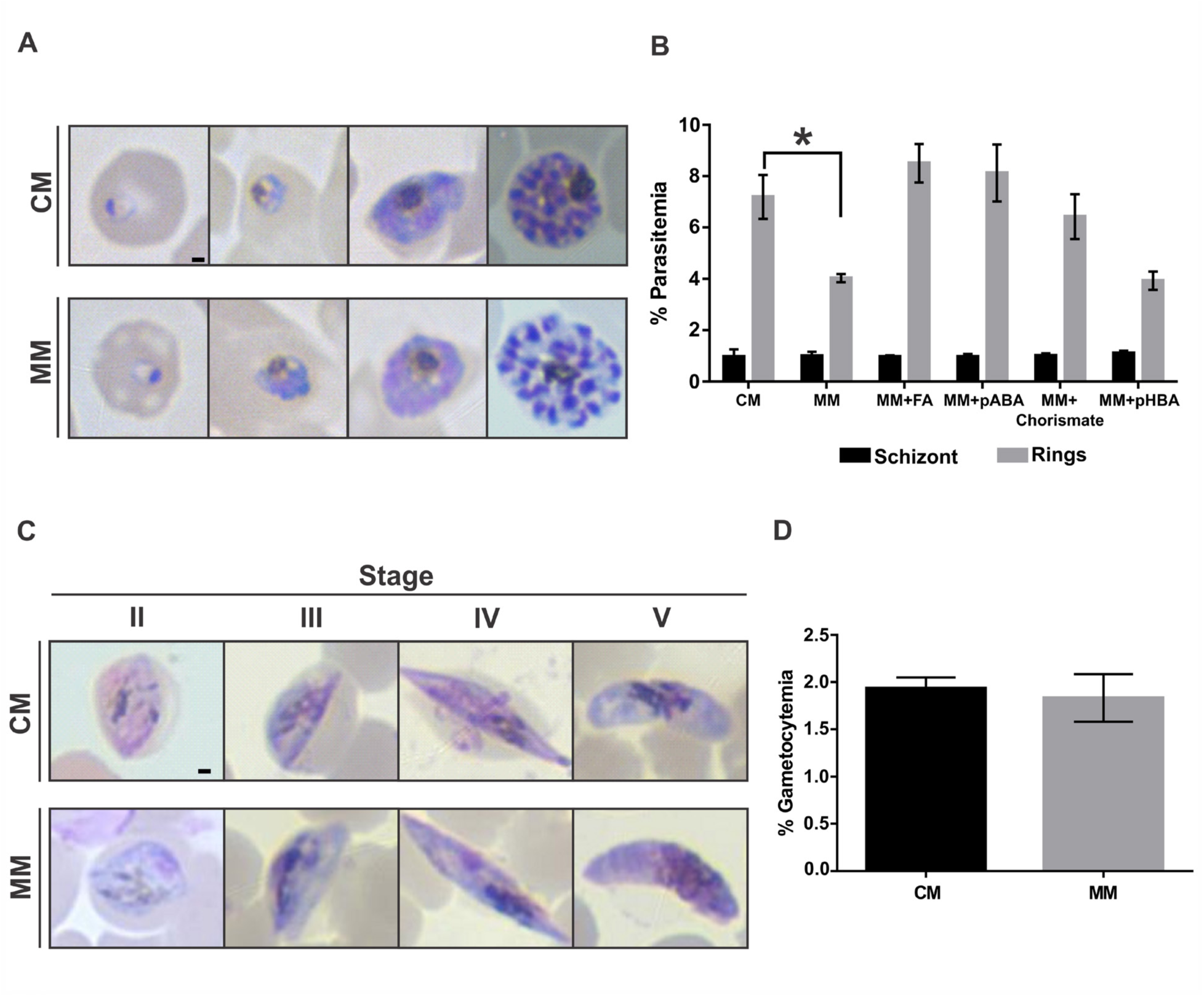
Development of *P. falciparum* asexual and gametocyte stages in the absence of exogenous supply of chorismate end-products. **(A)** Representative Giemsa-stained smears of *P. falciparum* asexual cycle of parasites grown in MM and CM. **(B)** Highly synchronous ring stage cultures from MM were supplemented with the indicated metabolites or CM and percentage of schizont stages after 24 h incubation and ring stages after 48 h incubation were assessed by Giemsa-stained smears and light microscopy. (*) indicates *p* < 0.001. **(C)** Representative Giemsa-stained smears of *P. falciparum* gametocyte stages obtained in MM and CM. **(D)** Final gametocytemia assessed by Giemsa-stained smears and light microscopy at day 12 after gametocytogenesis was induced in MM and CM. Scale bar indicates 1 μm. Values represent means ± S.E.M. of at least three independent assays.

Survival of early stage gametocytes is reduced by pyrimethamine, a well-known antifolate that targets dihydrofolate reductase (DHFR), supporting the importance of folate biosynthesis during gametocytogenesis ^29-31^. *P. falciparum* NF54 strain adapted to grow in MM was used to induce gametocytogenesis in parallel to *P. falciparum* NF54 strain grown in CM (see experimental procedures). As shown in Fig. 2C, gametocytes presented normal morphogenesis in the absence of exogenous supply of chorismate products. Differences in the timing of each stage during gametocytogenesis as well as final gametocytemia were not observed between parasites cultured in MM and CM (Fig. 2D). To our knowledge, this is the first time that gametocytogenesis in a pABA/FA-free medium is assessed where no differences were observed between CM and MM.

Altogether, our results support that both *P. falciparum* asexual and gametocyte stages exhibit normal development in MM, therefore, all further studies were performed using this defined medium.

### Reversal of growth inhibition to identify potential inhibitors of the shikimate pathway

After establishing the growth conditions in MM which allow the addition or replacement of metabolites in ^13^C-biolabeling experiments, we then assessed known inhibitors of the shikimate pathway as potential metabolic modulators that could be used also in gametocyte stages. Historically, metabolic inhibitors have been used to modulate metabolism and study biosynthesis of metabolites. Inhibitors are a valuable tool when genetic manipulation remains difficult to achieve or is not the best approach. This is the case for studying metabolism in *P. falciparum* gametocyte stages since continuous cultures lose the ability to generate gametocytes within a couple of months ^32^. Consequently, two previously reported potential inhibitors of the shikimate pathway, glyphosate ^7^ and (6*S*)-6-fluoroshikimate ^3^, were evaluated in parasites adapted to grow in MM. However, under our experimental conditions, glyphosate only partially inhibited parasite’s growth (∼40%) at 5 mM (Fig. S1) while 1 mM was previously reported to inhibit growth 100% 7. On the other hand, (6*S*)-6-fluoroshikimate was reported to inhibit *P. falciparum in vitro* growth up to 70% at 1 mM. We were able to synthesize (6*S*)-6-fluoroshikimate but under our experimental conditions only ∼30% inhibition was achieved at 1 mM (Fig. S1). Therefore, neither inhibitor was pursued as a potential metabolic modulator due to the high concentration required.

As an alternative to identify a more suitable inhibitor for our metabolic studies, a reversal of growth inhibition assay by end-products of chorismate was utilized to screen an open-access library from MMV named “Pathogen Box”, which contains a 2,4-diaminothieno [2,3-*d*-] pyrimidine derivative (MMV688345) previously reported as a potential antifolate in *P. berghei* and *P. gallinaceum* ^20^ as well as in *Toxoplasma gondii* ^21^. Since pHBA is not present in the standard RPMI medium but is postulated to be one of the end-products of chorismate, this metabolite was added to the screening medium at a final concentration of 7.3 μM. The pHBA concentration was selected based on the amount of pABA present in standard commercial RPMI (Table S1) as well as on the toxicity curve of pHBA alone (Fig. S2).

From the 400 compounds tested, only compound MMV688345 met our criteria of >90% growth inhibition at 5 and 1 μM for the compound alone and <60% growth inhibition in the presence of inhibitor and chorismate end-products, as detailed in the experimental procedures section. The complete data set listing growth inhibition for the 400 compounds tested is available as Supporting Information (DATA SET S1). MMV689437 also showed reduced inhibition in CM compared to MM, but it was not further pursued as a potential metabolic modulator due to its higher IC_50_ value (∼1000 nM) compared to MMV688345 (11.4 nM, Table 1) in the absence of chorismate end-products.

**Table 1.** Effect of exogenous metabolite supplementation on the inhibitory effect of MMV688345 against *P. falciparum* adapted to continuous culture in MM. Values represent the mean ± S.E.M. from at least three independent assays performed in triplicate. (*) Not significant by Benjamini and Hochberg procedure.

| Recovery Condition | IC<sub>50</sub> values (nM) | IC<sub>50</sub>-fold change vs MM | *p* value |
| --- | --- | --- | --- |
| CM | 1,162.0 ± 52.4 | 101.9 | 0.0006 |
| MM | 11.4 ± 0.8 | - |  |
| MM + shikimate (25 µM) | 30.7 ± 3.0 | 2.7 | 0.02 * |
| MM + shikimate-3-phosphate (25 µM) | 48.1 ± 6.1 | 4.2 | 0.03 * |
| MM + chorismate (12.5 µM) | 964.8 ± 56.9 | 84.6 | 0.0005 |
| MM + pABA (7.3 µM) | 910.8 ± 54.2 | 80 | 0.00007 |
| MM + FA (7.3 µM) | 553.5 ± 21.1 | 48.5 | 0.00001 |
| MM + tyrosine (111 µM)<br>phenylalanine (90 µM)<br>tryptophan (24.5 µM) | 43.1 ± 5.8 | 3.7 | 0.01 * |
| MM + pHBA (7.3 µM) | 49.6 ± 5.5 | 4.3 | 0.01 * |

MMV688345 was retested in a dose-dependent manner in MM and CM (Fig. 3), and in MM containing each individual chorismate end-product present in the screening CM. Recovery of growth inhibition experiments showed that neither pHBA nor the mix of aromatic amino acids was able to significantly increase the IC_50_ values (Table 1), while pABA and FA shifted the IC_50_ values over 80- and 48.5-fold respectively, demonstrating that these two metabolites are necessary for parasite survival in the presence of MMV688345 pressure.

**Figure 3.**
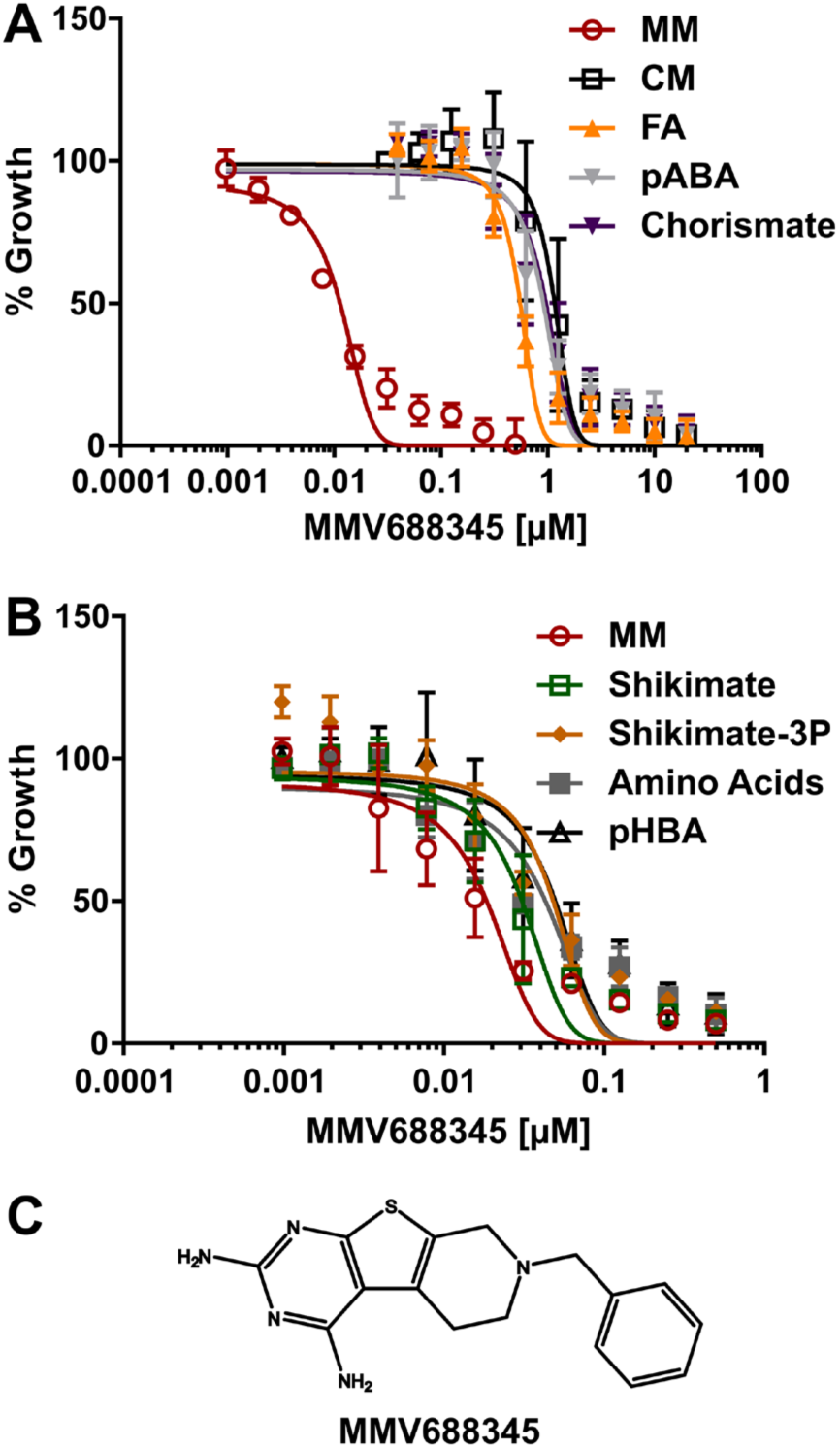
MMV688345 dose-dependent growth inhibition and growth recovery by metabolite supplementation. **(A)** MMV688345 dose-dependent reversal of growth inhibition by CM, FA (2.2 μM), pABA (7.3 μM), and chorismate (12.5 μM). **(B)** MMV688345 dose-dependent growth inhibition was not reversed by shikimate (25 μM), shikimate 3-phosphate (25 μM), aromatic amino acids (tyrosine at 111 μM, phenylalanine at 90 μM, and tryptophan at 24.5 μM), or pHBA (7.3 μM). **(C)** MMV688345 chemical structure.

Interestingly, chorismate alone also shifted the IC_50_ values over 84.6-fold while shikimate-3-phosphate and shikimate did not significantly shift the IC_50_ values (Fig. 3 and Table 1), despite uptake of these intermediates by the malaria parasite since both metabolites increased growth of *P. falciparum* when added to MM (Fig. S3). The results suggest that the potential molecular target of MMV688345 is among the last two steps of chorismate biosynthesis (Fig. 1) and that the major metabolic role of chorismate in the malaria parasite is to supply folate precursors. Thus, we further tested this hypothesis by metabolomics.

It is worth mentioning that the IC_50_ values reported in Table 1 were reproducible using different medium preparations and blood batches during the course of our investigation. Uninfected erythrocytes were washed and stored in MM containing Albumax II as these cells do not have a known requirement for folate. During erythropoiesis, erythrocytes accumulate 5-methyl tetrahydrofolate since mature erythrocytes lack a functional γ-polyglutamate hydrolase. The 5-methyl tetrahydrofolate remains bound to hemoglobin due to its high charge, thus, unavailable to the malaria parasite ^14^. Moreover, pABA is the only known metabolite present in adult human plasma at variable concentrations (0.15 - 6.56 μM) ^26^. Thus, potential differences in pABA concentrations present in the plasma of the anonymous blood donors could result in different concentration of pABA inside of erythrocytes, thus, resulting in variable IC_50_ values between blood batches.

### MMV688345 is cytocidal in schizont stage and does not affect late stage gametocytes

Before performing metabolomics analysis, we first assessed if MMV688345 was cytocidal and parasite stage-specific by determining the 50% lethal concentration (LC_50_) at 72 h using highly synchronous cultures starting at ring stage and bolus incubation times ranging from 24 to 48 h. At each indicated time, MMV688345 was washed out and parasites were returned to culture to complete 72 h growth, at which point the growth curves were measured by SYBR Green assay (Fig. 4A). In parallel, stage assessment of parasites was performed by Giemsa-stained thin smears of MMV688345-treated and control parasites at the time that MMV688345 was washed out as well as after completing 72 h growth (Fig. 4B). The SYBR Green assay at 72 h confirmed that parasites were still able to grow and reinvade erythrocytes following 24 h incubation with MMV688345 since rings were observed after 48 h incubation in the absence of the inhibitor and the LC_50_ values were higher than the IC_50_ values at 72 h performed simultaneously with these assays (28 ± 5 nM). However, when incubation with MMV688345 was extended for 36 and 48 h, the LC_50_ values were 50 ± 11 nM and 24 ± 4 nM, respectively, which were very close to the measured IC_50_ value. The cytocidal action of MMV688345, therefore, reveals at 36 h, which is the time needed for parasites to progress to the schizont stage where the need for precursors of folate such as pABA also increases to sustain DNA replication ^33-35^.

**Figure 4.**
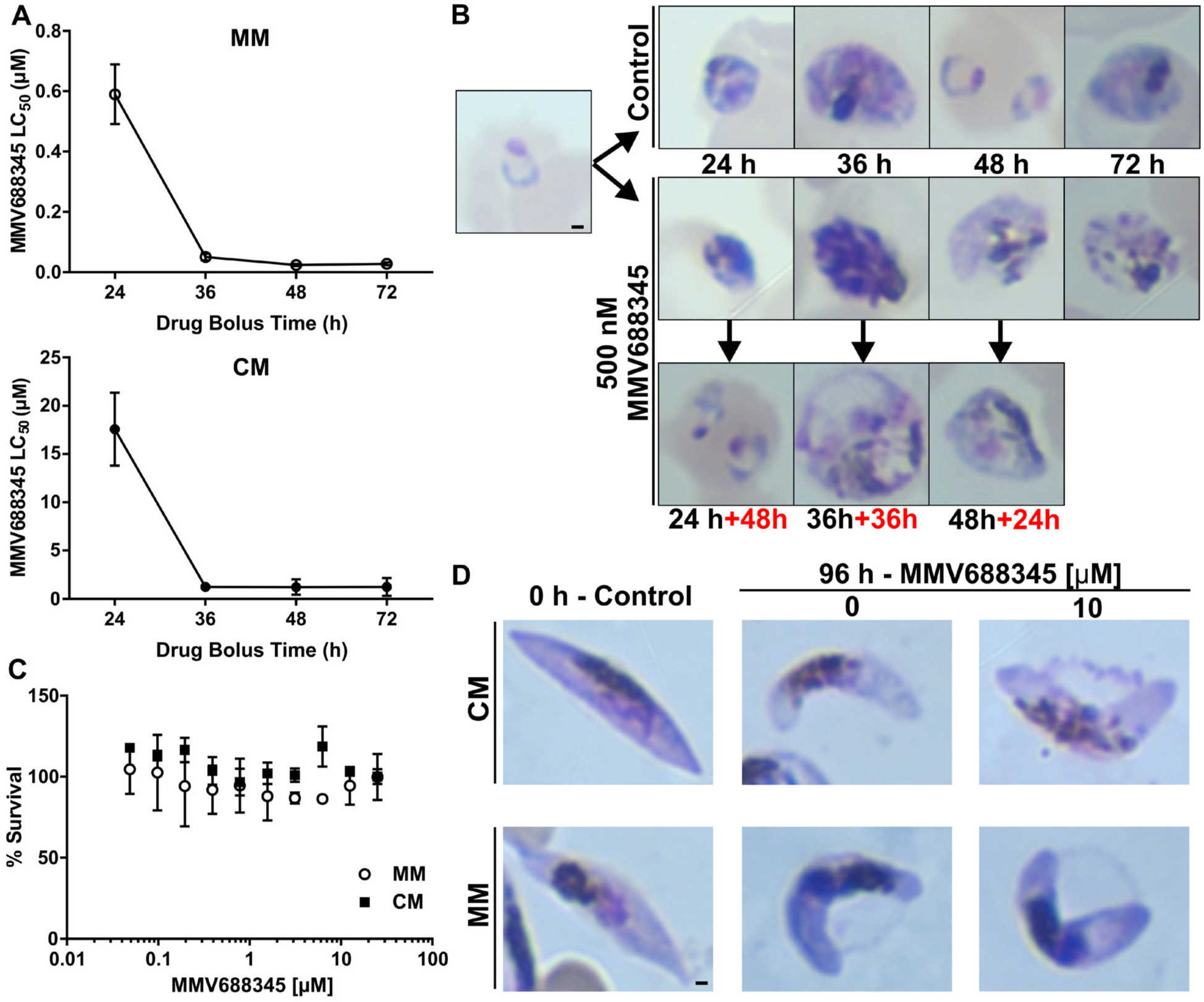
Measured MMV688345 LC_50_ decreases with increasing drug exposure time. **(A)** MMV688345 becomes cytocidal at 36 h drug bolus time when parasites reach schizont stages both in MM and CM. **(B)** Giemsa-stained thin smears of MMV688345-treated and control parasites at the time that MMV688345 was washed out as well as after completing 72 h of growth (added time in red). (**C)** Survival of late stage gametocytes was assessed by AlamarBlue both in MM and CM. **(D)** Giemsa-stained thin smears of MMV688345-treated and control late-stage gametocytes.

Consistent with a potential antifolate mechanism of action ^20,21^, MMV688345 was not active against late stage gametocytes (Fig. 4C-D) as the folate-mediated pyrimidine synthesis required for DNA replication is not required until male gametocyte exflagellation ^31,36^.

### MMV688345 does not affect ubiquinone biosynthesis

As mentioned above, the current available biochemical and genetic information suggest that the potential major metabolic role of chorismate in the malaria parasite is to supply folate precursors during the intraerythrocytic cycle ^11-13^. Thus, to further investigate the metabolic connection of the shikimate pathway to ubiquinone biosynthesis a metabolomic analysis was performed comparing isoprenoid products of MMV688345-treated and untreated cultures. Highly synchronous cultures at ring stage were treated with 500 nM MMV688345 in parallel with untreated control cultures. Uninfected erythrocytes were also treated as a control. After 20 h of treatment, *P. falciparum* cultures were smeared to verify that both untreated and treated cultures were at the schizont stage and that they did not present morphological defects due to the treatment which may confound the metabolomics results (Fig. 5A). As a control, a sample from untreated and MMV688345-treated parasites were further cultured to complete 48 h of treatment when morphological defects became evident due to the treatment. We performed a principal component analysis (PCA) where results are displayed as score plots, and each point represents a sample that when clustered together indicates similar metabolite composition based on the untargeted analysis performed (Fig. 5B). This analysis revealed that both untreated and MMV688345-treated groups have similar isoprenoid product composition and further analysis confirmed that ubiquinone-9 levels, the major ubiquinone species detected in *P. falciparum* by LC-HRMS, were not affected by MMV688345 treatment (Fig. 5C). Previous studies using radiolabeled metabolic precursors showed that ubiquinone biosynthesis occurs throughout the intraerythrocytic cycle and increases at the end of the intraerythrocytic cycle when the parasite is preparing for schizogony ^15,37^.

**Figure 5.**
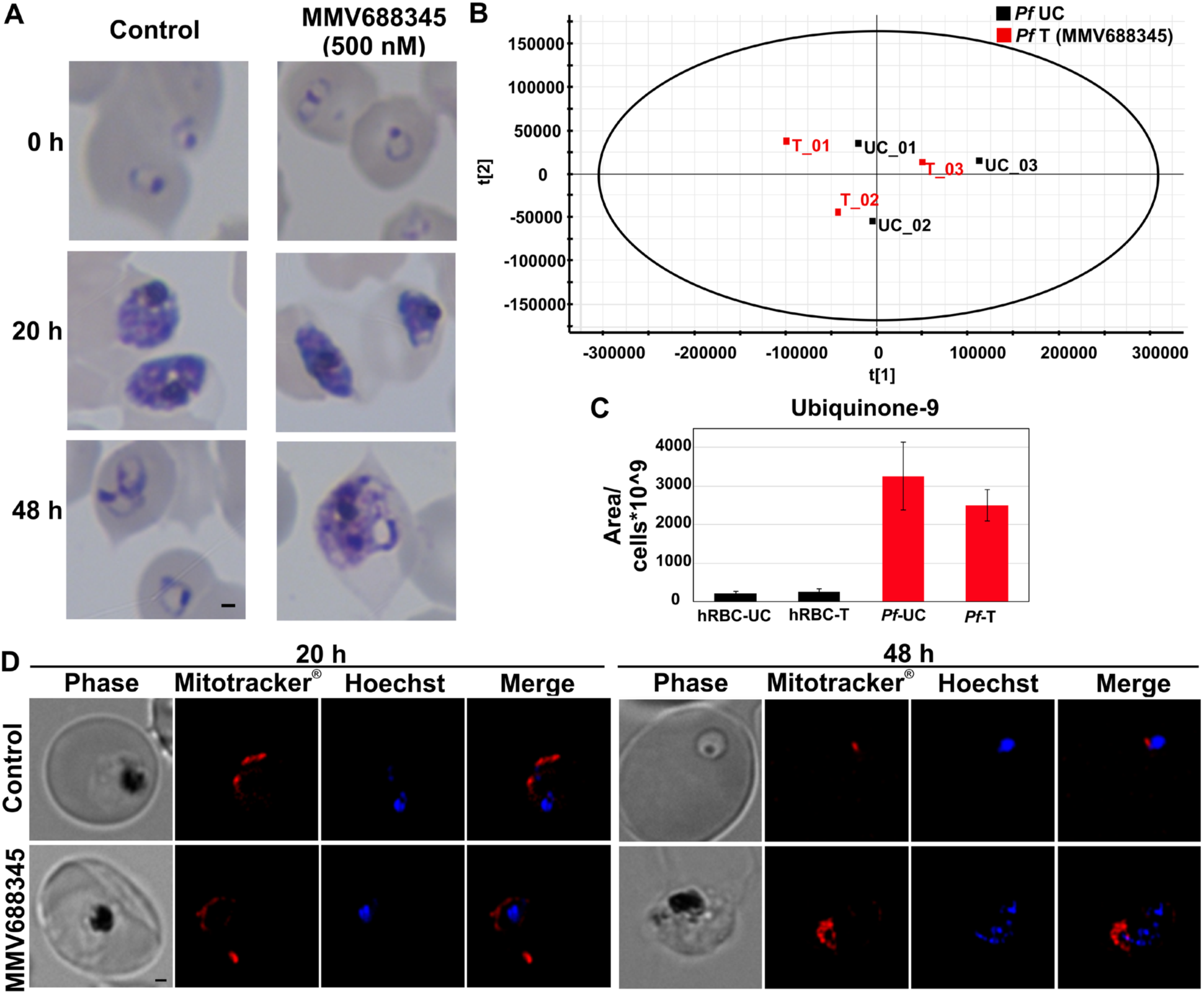
MMV688345 does not affect ubiquinone biosynthesis. **(A)** Giemsa-stained thin smears of MMV688345-treated and control parasites. After 20 h of treatment, *P. falciparum* cultures were recovered for metabolomics analysis and a sample of each condition was maintained in culture to complete 48 h of treatment. **(B)** PCA model score plot of LC-HRMS data from analyzed *P. falciparum* samples comparing MMV688345-treated (*Pf*-T) and untreated control (*Pf*-UC) samples. **(C)** Levels of ubiquinone-9 detected in *P. falciparum* by LC-HRMS were not affected by MMV688345 treatment. **(D)** Effect of MMV688345 treatment on the mitochondrial membrane potential was assessed using MitoTracker^®^ after 20 and 48 h of treatment.

Ubiquinones play a key role as electron acceptors in the mitochondrial electron transport chain ^38^. Thus, we assessed the changes in mitochondrial membrane potential using MitoTracker^®^ as further confirmation that MMV688345 treatment does not affect ubiquinone biosynthesis. As shown in Fig. 5D, mitochondrial function was not affected by MMV688345 treatment even after 48 h in the presence of inhibitor when parasites are morphologically affected by MMV688345 treatment (Fig. 5A).

### Lack of metabolic connection between chorismate and ubiquinone biosynthesis

Evidence of a potential lack of metabolic connection between chorismate and ubiquinone biosynthesis was first suggested, although not explicitly, by Roberts and colleagues where *P. falciparum* growth inhibition by glyphosate was abolished in the presence of only pABA or FA ^7^. Shortly after, McConkey reported a similar result using (6*S*)-6-fluoroshikimate ^3^. Moreover, no genes can be found for the first enzyme of ubiquinone biosynthesis, chorismate lyase (EC 4.1.3.40), which converts chorismate into pHBA or for the enzyme responsible for *C*1-decarboxylation of the head group which has only been identified in prokaryotes ^39^. In addition, recent studies suggest that chorismate synthase is not essential for *Plasmodium* survival ^11,13^ and our results suggest that the head group of ubiquinones in *P. falciparum* may not derive from chorismate or that an alternative source of pHBA may exist. It has been reported that pHBA in human plasma is around 2.5 μM ^40^ and that it is taken up by human erythrocytes ^41^, despite the lack of ubiquinone biosynthesis in this cell. Therefore, it is possible that asexual intraerythrocytic stages rely on the host supply for synthesis of ubiquinones. Interestingly, yeast can also use pABA as an alternative precursor for the head group ^42,43^. Thus, we tested whether *P. falciparum* is able to use [^13^C_6_]pHBA and/or [^13^C_6_]pABA as metabolic precursors for ubiquinone biosynthesis. Parasites were pretreated with 500 nM MMV688345 to reduce endogenous unlabeled pABA and potentially pHBA before adding the ^13^C-biolabeling precursors and samples were analyzed by LC-HRMS as described in the experimental procedures. Surprisingly, neither [^13^C_6_]pHBA nor [^13^C_6_]pABA was incorporated into ubiquinone-9 (Fig. S4) in all three independent experiments under the conditions tested.

Ubiquinone biosynthesis is active in the malaria parasite ^15,16^, and it is postulated to be essential in *P. falciparum* asexual stages ^12,13^, however, it remains to be experimentally confirmed. Altogether, our results suggest that it is possible the benzoquinone ring of ubiquinone in *P. falciparum* may be synthesized through an unidentified route.

### Screening using reversal of growth inhibition by FA and pABA may favor identification of competitive antifolates

The lack of evidence connecting chorismate and the biosynthesis of the benzoquinone ring suggested by our metabolomics analysis led us to revise our hypothesis that the potential molecular target of MMV688345 could be among the last two steps of chorismate biosynthesis. In a previous report of the open-source compound library from MMV called “Malaria Box”, a folate pathway inhibition assay was used by Gamo and colleagues to aid in the identification of the mechanism of action of these compounds ^24^. They adapted a *P. falciparum* 3D7A strain to grow in low levels of FA (0.227 μM) in a RPMI medium depleted of pABA and folinic acid. In their analysis, the presence of three potential antifolate compounds (MMV667486, MMV667487 and MMV396678) could be identified ^24^. MMV667486 and MMV667487 are analogs of cycloguanil that targets DHFR. A previous metabolomics study that aimed to aid in the identification of the mechanism of action of compounds among the “Malaria Box” has also indicated that MMV667487 may target folate biosynthesis ^44^. We tested these three candidates under our experimental conditions described above using pyrimethamine as a control. As expected, growth inhibition observed in MM by pyrimethamine (IC_50_ = 0.18 ± 0.03 nM), MMV667486 (IC_50_ = 1.28 ± 0.17 nM) and MMV667487 (IC_50_ = 0.71 ± 0.05 nM) was reversed in CM while growth inhibition by MMV396678 (IC_50_ = 8,723 ± 1.3 nM) was not reversed by FA (Fig. S5). Surprisingly, chorismate, FA and pABA but not shikimate, also reversed growth inhibition by pyrimethamine, MMV667486 and MMV667487 similar to MMV688345 (Fig. S5).

To our knowledge, this is the first time that chorismate has been shown to rescue growth inhibition of antifolates in the malaria parasite. Pyrimethamine is a known competitive inhibitor of DHFR; thus, the presence of high concentrations of FA and pABA are expected to outcompete pyrimethamine from its enzyme target ^45^, while chorismate may increase the synthesis of pABA and dihydrofolate, therefore, competing with the inhibitor. A similar response was observed with MMV688345, MMV667486 and MMV667487, suggesting that the molecular target localizes downstream chorismate and not within the shikimate pathway as we initially hypothesized. However, it is puzzling that neither shikimate nor shikimate-3-phosphate were able to significantly reverse growth inhibition of any of the antifolates tested despite uptake of these intermediates by the malaria parasite since both metabolites increased growth of *P. falciparum* when added to MM (Fig. S3).

Reversal of growth inhibition or metabolic suppression by metabolites has already aided in identifying the mechanism of action of biologically active molecules including the *P. falciparum* IspD inhibitor MMV008138 ^46,47^ as well as new antibacterial inhibitors ^48^. However, due to the potential role of chorismate as a precursor in *de novo* folate biosynthesis and the complexity of folate metabolism in the malaria parasite where the contribution of the *de novo* biosynthesis and salvage pathways is starting to be elucidated ^49,50^, the use of reversal of growth inhibition will likely result in the selection of competitive antifolates even when chorismate is used as a metabolic suppressor of growth inhibition ^24^. Thus, the use of *in vitro* enzymatic reactions as screening platforms using recombinant proteins of selected targets may be a better approach to identify or rationally design new non-competitive and specific antifolates ^51^. Folate metabolism remains one of the best established clinical targets due to its safety for administration to infants, children and pregnant women, the most vulnerable populations affected by malaria ^52^.

## Conclusion

In summary, we systematically assessed the development of both asexual and sexual stages of *P. falciparum* in a defined medium in the absence of an exogenous supply of chorismate end-products that will aid future research to elucidate the contribution of *de novo* biosynthesis and the salvage pathway to folate metabolism in *P. falciparum*. We utilized reversal of growth inhibition by different intermediates of the shikimate pathway and chorismate products to study the mechanism of action of previously identified antifolates which suggested that a different approach may be required to avoid selection of competitive inhibitors. Lastly, we presented biochemical evidence suggesting that a different pathway may be used by the malaria parasite to synthesize the benzoquinone which guarantees further investigation.

## Materials and Methods

### Parasite cultures

*P. falciparum* NF54 (sulfadoxine-resistant) strain was obtained through the MR4 Malaria Reagent Repository (ATCC, Manassas, VA) as part of the BEI Resources Repository, NIAID, NIH. Parasites were adapted to grow in RPMI Minimal Media (MM) without Phenol Red, pABA, FA, tryptophan, phenylalanine and tyrosine which are present in the standard RPMI media (CM) (Table S1). Parasites were maintained at 4% hematocrit in O^+^ human erythrocytes (The Interstate Companies, TN) in MM supplemented as indicated in Table S1. Cultures were maintained at 37°C under reduced oxygen conditions (5% CO_2_, 5% O_2_, and 90% N_2_). Supplementation of the MM with individual metabolites or in combination is indicated in the results section. Standard RPMI media (CM) used in this work was obtained by supplementing MM with pABA, FA, tryptophan, phenylalanine and tyrosine at the concentrations reported for Gibco™ RPMI 1640 Medium (powder). Supplementation with 7.3 μM pHBA was performed as indicated in the result section.

### Assessment of gametocytogenesis in MM and with MMV688345 treatment

Gametocyte stages were obtained using *P. falciparum* NF54 strain revived in MM and adapted for two weeks before inducing gametocytogenesis. In parallel, gametocytes were obtained in CM to assess potential differences in gametocyte development. Briefly, highly synchronous cultures at young ring stage were obtained by 5% sorbitol treatment and purified using a MACS^®^ magnetic affinity column (day 0) and set at 8-12% parasitemia and 5% hematocrit. On day 0 and 1, gametocytogenesis was induced by incubating cultures with 50% spent media and 50% fresh media for 40-48 h, as described previously ^53^. After reinvasion, asexual parasites were depleted from culture by addition of fresh media containing 20 U/mL of heparin for 4 days (day 2 to 5) and sorbitol treatment ^54^. After day 5, the media was replaced every two days until gametocytes reached stage V. Development of parasites was monitored by microscopic evaluation of Giemsa-stained thin smears.

In order to assess whether MMV688345 affects late stage gametocytes, stage IV gametocytes were generated as described above and incubated for 72 h in the presence of increasing concentrations of the inhibitor (10-points curve). AlamarBlue was then added at 10% of the well volume as previously described and incubated for another 24 h ^46,55^. In addition, untreated controls and treated gametocytes with 10 μM MMV688345 were morphologically assessed by Giemsa-stained thin smears as shown in Fig. 4.

### Chemicals

The Malaria Box and Pathogen Box compound collections, containing 400 compounds in each library, were kindly provided by Medicines for Malaria Venture (MMV, Switzerland) and supplied at 10 mM in dimethyl sulfoxide (DMSO). The original samples were aliquoted to individual library sets with a final drug concentration of 1 mM in DMSO, as recommended by MMV and stored at −80°C. Resupply of MMV688345 was provided by Evotec (US) Inc. for MMV. The complete information regarding the Pathogen Box and Malaria Box composition can be found at https://www.mmv.org. Synthesis of (6*S*)-6-fluoroshikimic acid was performed according to the literature procedure ^56^.

### Reversal of growth inhibition by chorismate end-products

We assessed whether compounds from the Pathogen Box were targeting chorismate biosynthesis by measuring growth recovery in the presence of inhibitor and 2.2 μM FA, 7.3 μM pABA, 7.3 μM pHBA, 111 μM tyrosine, 24.5 μM tryptophan and 90 μM phenylalanine. Briefly, *P. falciparum* ring stages were cultured under the following conditions: control (no drug), 5 μM or 1 μM of Pathogen Box compounds, and 5 μM or 1 μM of Pathogen Box compounds with chorismate end-products indicated above. Artemisinin was used as a control drug for growth inhibition as it is not affected by presence or absence of chorismate end-products. All conditions were set in 96-well half-area dark plates (100 µL/well with 1% hematocrit and 1% parasitemia), incubated for 72 h, and analyzed by the SYBR green assay as described previously ^57,58^. Percentage of growth was normalized to untreated control parasites. Since several compounds within the Pathogen Box did not inhibit growth completely at 5 or 1 μM, we only considered reversal of growth inhibition when compounds alone inhibited growth >90% at 5 and 1 μM and <60% in the presence of inhibitor and chorismate end-products. Uninfected erythrocytes were used for background determination.

In order to rule out potential toxicity from each metabolite used in this study, a ten-point curve of increasing concentrations of each metabolite was performed (Fig. S2). To establish dose-dependency for the reversal of growth inhibition, *P. falciparum* ring stage parasite cultures (100 µL/well, 1% hematocrit, and 1% parasitemia) were grown for 72 h in the presence of increasing concentrations of the inhibitor (10-points curve) and in the presence or absence of metabolites at fixed concentrations as indicated in the results section. Parasite growth was assessed by SYBR green as described previously ^57,58^. The half maximal inhibitory concentrations (IC_50_) were calculated from dose-response curves calculated from normalized percent activity values from three or more independent determinations performed in triplicate and reported as means ± S.E.M. The IC_50_ values were determined from log10-transformed concentrations using GraphPad Prism 6 software (GraphPad Software, Inc.) and nonlinear regression curve fitting.

### Cytocidal (LC_50_) assay

In order to determine the concentration of a bolus dose of MMV688345 that kills 50% of parasites (LC_50_), *P. falciparum* adapted to MM cultures were exposed to increasing concentrations of MMV688345. The drug was subsequently washed away at 24, 36, and 48 h to probe stage specificity for the activity of MMV688345 as described previously ^59,60^. Briefly, following the bolus dose incubation, plates were centrifuged at 700 *x g* for 3 min and MMV688345-containing medium was removed. Cell pellets were washed three times with 100 μl of MM and re-suspended in 100 μl of MM without the inhibitor. Washed plates were then incubated at 37 °C to complete a total of 72 h after setting the assays, and growth of surviving parasites was assessed by SYBR Green. The half-maximum lethal concentration (LC_50_) values were calculated with GraphPad Prism 6 software (GraphPad Software, Inc.) using nonlinear regression curve fitting. The reported values represent means ± S.E.M. of at least three independent assays, with each assay performed in triplicate. Parasites untreated or treated with MMV688345 at the IC_100_ concentration (500 nM) were smeared and stained with Giemsa before washing the inhibitor in order to assess stage development at the time that MMV688345 was removed as well as at the end of the experiment (72 h) as shown in Fig. 4.

### Statistical analysis

The *t*-test and Benjamini and Hochberg procedure were used to analyze differences between different conditions, and a false discovery rate of 0.01 was used to identify statistically significant differences. The principal component analysis (PCA) was performed using Progenesis QI software (Nonlinear Dynamics, Waters Corporation, Milford, MA, USA).

### Sample preparation for profiling of isoprenoids

Mycoplasma-free parasite cultures were tightly synchronized by 5% sorbitol treatment. Each biological replicate was obtained from three 75 cm^2^ flasks of *P. falciparum* cultures in MM at 4% hematocrit, starting with early *P. falciparum* ring stages at 7% parasitemia and harvesting after 20 h incubation under different conditions at the schizont stage. In order to assess the effect of MMV688345 on downstream isoprenoids, untreated or 500 nM MMV688345-treated *P. falciparum* cultures were incubated for 20 h in parallel with untreated and treated uninfected erythrocytes as control. For metabolic labeling experiments using [^13^C_6_]pHBA (7.3 μM) and [^13^C_6_]pABA (7.3 μM) as potential metabolic precursor for ubiquinone biosynthesis, highly synchronous ring-stage cultures were pretreated with 500 nM MMV688345 for 6 h and ^13^C-biolabeling precursors were added directly into MM and incubated for 14 h in the presence of the inhibitor. All ^13^C-biolabeling precursors were purchased from Cambridge Isotope Laboratories (CIL). In all cases, infected erythrocytes were released from the host cell by lysis with 0.03% saponin in cold PBS containing 2 g/L glucose and washed three times with PBS/glucose by centrifuging 7 min at 10,000 *× g* at 4°C and 10 μL were collected in the last wash to count parasites using a Countess cell counter (Thermo Fisher). Cell pellets were flash-frozen and stored at −80°C until the metabolite extraction which was performed by treating samples with 1 mL cold methanol and three consecutive extractions with 2 mL of hexane followed by pulse-vortex for 1 min and 10 min sonication in an ultrasonic bath. The upper phases were combined, dried under nitrogen and stored at −80°C until LC-HRMS analysis. For LC-HRMS analysis, samples were resuspended in 100 µL of methanol/acetonitrile/isopropanol (60:25:15, v/v) pulse vortexed 5 times and sonicated for 10 min. Samples were protected from light during the entire procedure.

### Ultra-performance liquid chromatography and high-resolution mass spectrometry analysis (LC-HRMS)

Isoprenoid analysis was performed on an IonKey/MS system composed of an ACQUITY UPLC M-Class, the ionKey source, and an iKey HSS T3, 100 Å, 1.8 µm (particle size), 150 µm × 100 mm column coupled to a Synapt G2-Si mass spectrometer (Waters Corporation, Milford, MA, USA). Isoprenoid separation was accomplished using a binary gradient system consisting of methanol/acetonitrile (3:1, v/v), with 10 mM ammonium formate + 0.1% formic acid (mobile phase A) and isopropanol/acetonitrile (3:1, v/v) containing 10 mM ammonium formate + 0.1% formic acid (mobile phase B). Sample analysis was performed using a linear gradient over a total run time of 35 min. The gradient was programmed as follows: 0.0-2.0 min 18% B, 2-16 min from 18% B to 89% B, 16-20 min at 89 % B, 20-25 min from 89% B to 18% B, and re-equilibrated for 10 min. The flow rate was 3 µL/min, the injection volume was 5 µL in full loop mode, and the iKey temperature was set at 40°C. MS analyses were performed acquiring in MS^E^ mode from 80 to 1500 *m/z* in positive electrospray ionization mode with a capillary voltage of 3 kV. Data were collected in two channels throughout the entire acquisition: low collision energy (6.0 V) for the molecular ions and high collision energy (15-40 V) for product ions. The source temperature was set at 110°C. Leucine enkephalin (50 pg/mL) was used as the lock mass (m/z 556.2771) with parameters set to 1 s scan at 20 s intervals. Infusion flow rate for lock mass was 1 µL/min. For unlabeled samples, peak identification, alignment, normalization, and statistical treatment of the data was performed using Progenesis QI software (Nonlinear Dynamics, Waters Corporation, Milford, MA, USA). For ^13^C-biolabeled samples, MassLynx software (Waters Corporation, Milford, MA, USA) was used for data processing and isotopomer labeling distribution analysis.

### Fluorescence microscopy

A DeltaVision Microscope System I with a 100x/1.4NA oil immersion objective. CoolSnap HQ2 high resolution CCD camera and the acquisition software SoftWorx were used to image live parasites as described previously with minor modifications ^46^. Briefly, parasites were treated with 500 nM MMV688345. Aliquots from untreated control and treated cultures were obtained at 20 and 48 h after treatment started and incubated with 100 nM MitoTracker^®^-Red-CM-H2XRos (Life Technologies) for 15 min at 37°C. Parasites were washed once with MM and 0.25 µg/mL Hoechst 33342 (Life Technologies) was added to each condition immediately before imaging. Final figures were prepared using Adobe Photoshop CS2.

## Acknowledgements

This work was supported by the National Institutes of Health (AI108819 to M.B.C.). We thank Medicines for Malaria Venture (MMV, Switzerland) for supply of the Pathogen Box, Malaria Box and MMV688345. The following reagent was obtained through MR4 as part of the BEI Resources Repository, NIAID, NIH: NF54, MRA-1000 by M. Dowler, Walter Reed Army Institute of research.

## Author Contributions

A.L.V., M.L.F-M, E.F.M., M.B.C. conceived of and designed the work. A.L.V., M.L.F-M, E.F.M., N. H., G.J.B. performed data acquisition, analysis and validation. K.J.S., P.C.T. performed chemical synthesis. A.L.V., E.F.M. and M.B.C wrote the original draft. All authors reviewed and edited the text.

## Competing interests

The authors declare no competing interests.

