## Supplemental Information for "Metabolic dependency of chorismate in *Plasmodium falciparum*"

Running title: *Chorismate dependency in P. falciparum*

**Table S1.** RPMI minimal and complete composition used in this study.

| Nutrient | Final Concentration (µM or as indicated) | Minimal Medium (MM) | Complete RPMI Medium (CM) |
|---|---|---|---|
| L-Arginine | 200 | + | + |
| L-Asparagine | 200 | + | + |
| L-Aspartic acid | 150 | + | + |
| L-Cysteine | 200 | + | + |
| L-Glutamic acid | 136 | + | + |
| L-Glutamine | 200 | + | + |
| Glycine | 133 | + | + |
| L-Histidine | 97 | + | + |
| Hydroxy-L-proline | 153 | + | + |
| L-Isoleucine | 100 | + | + |
| L-Leucine | 381 | + | + |
| L-Lysine | 200 | + | + |
| L-Methionine | 101 | + | + |
| L-Proline | 174 | + | + |
| L-Serine | 200 | + | + |
| L-Threonine | 168 | + | + |
| L-Valine | 171 | + | + |
| Choline Chloride | 0.0214 | + | + |
| D-Biotin | 0.00082 | + | + |
| D-Calcium pantothenate | 0.00052 | + | + |
| *Myo*-Inositol | 0.194 | + | + |
| Niacinamide | 0.0081 | + | + |
| Pyridoxine hydrochloride | 0.0048 | + | + |
| Riboflavin | 0.00052 | + | + |
| Thiamine hydrochloride | 0.00296 | + | + |
| Vitamin B12 | 0.00037 | + | + |
| Reduced Glutathione | 3,254 | + | + |
| $Ca(NO_3)_2$ | 609 | + | + |
| HEPES | 20,967 | + | + |
| KCl | 5,365 | + | + |
| $MgSO_4$ | 407 | + | + |
| NaCl | 102,669 | + | + |
| $Na_2HPO_4 \cdot 7\ H_2O$ | 2,797 | + | + |
| D-Glucose | 22,203 | + | + |
| $NaHCO_3$ | 26,784 | + | + |
| Gentamicin | 41,876 | + | + |
| Hypoxanthine | 367 | + | + |
| Albumax II | 5 (g/L) | + | + |
| **Folic Acid** | **2.2** | | + |
| ***p*-Aminobenzoate** | **7.3** | | + |
| **L-Phenylalanine** | **90** | | + |
| **L-Tryptophan** | **24.5** | | + |
| **L-Tyrosine** | **111** | | + |
| ***p*-Hydroxybenzoate** | **7.3** | | + |

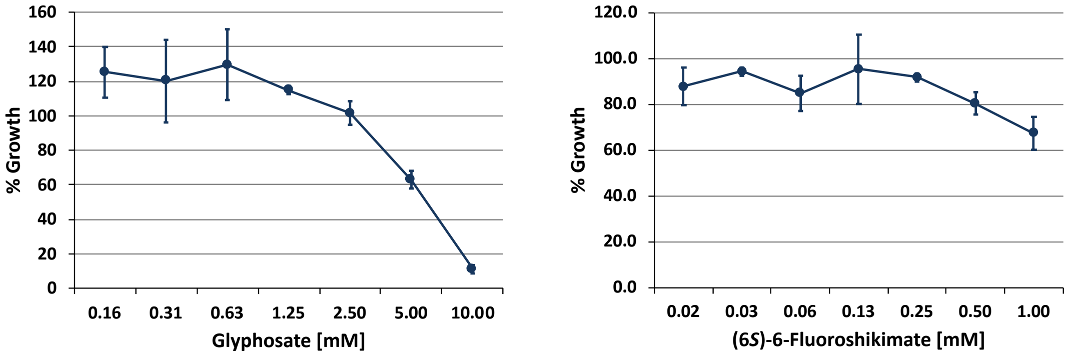


**Fig. S1. Effect of glyphosate and (6*S*)-6-fluoroshikimate on *in vitro* growth of *P. falciparum*.** Dose-dependent growth inhibition was determined after incubation for 72 h in the presence of increasing concentrations of the inhibitor in MM. Parasite growth was assessed by SYBR green. Results represent means ± S.E.M. of two independent assays, with each assay performed in triplicate.

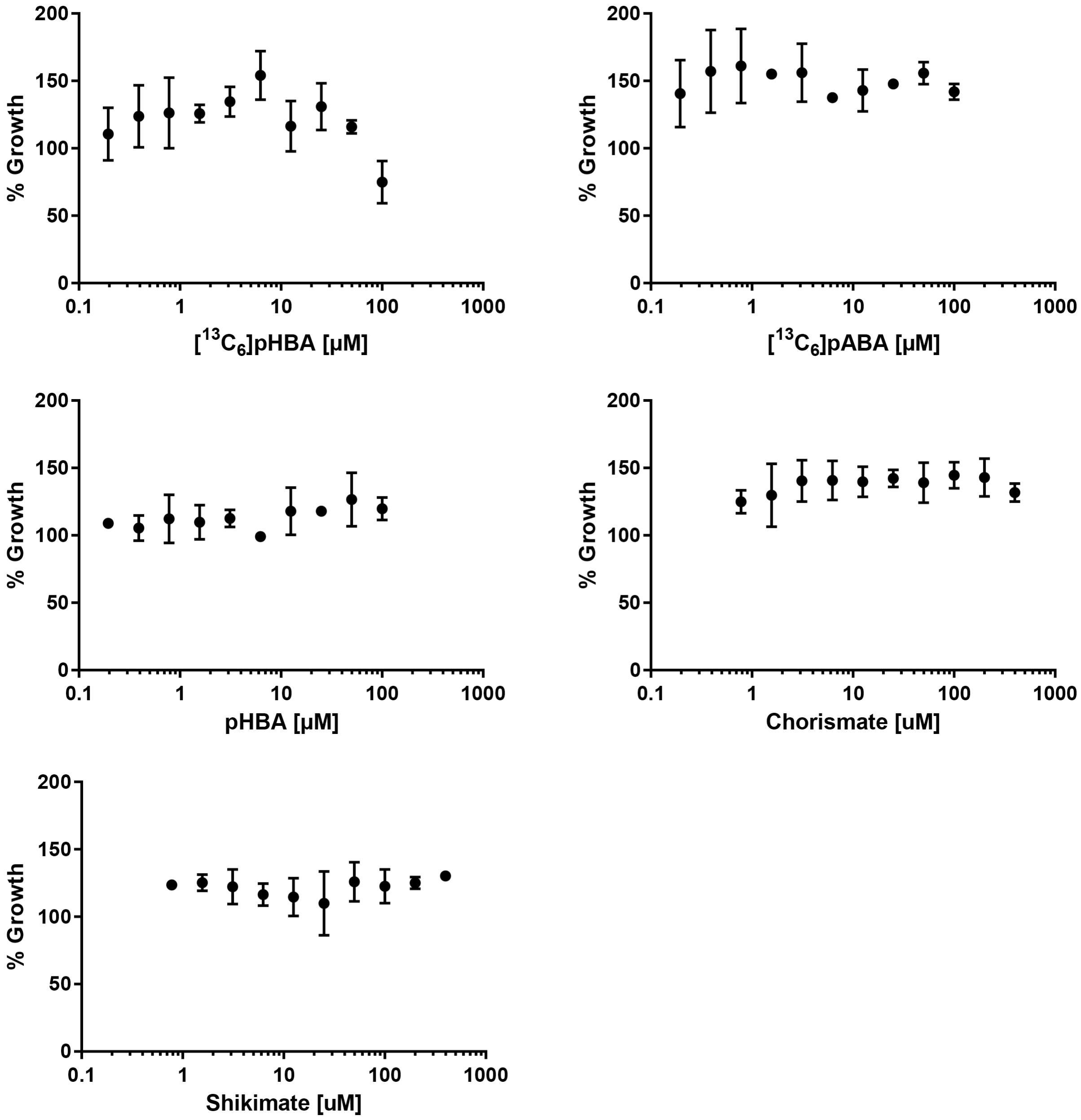


**Fig. S2. Effect of metabolites on *P. falciparum in vitro* growth.** Concentration-dependent potential metabolite toxicity was assessed in MM by SYBR green assay after 72 h incubation.

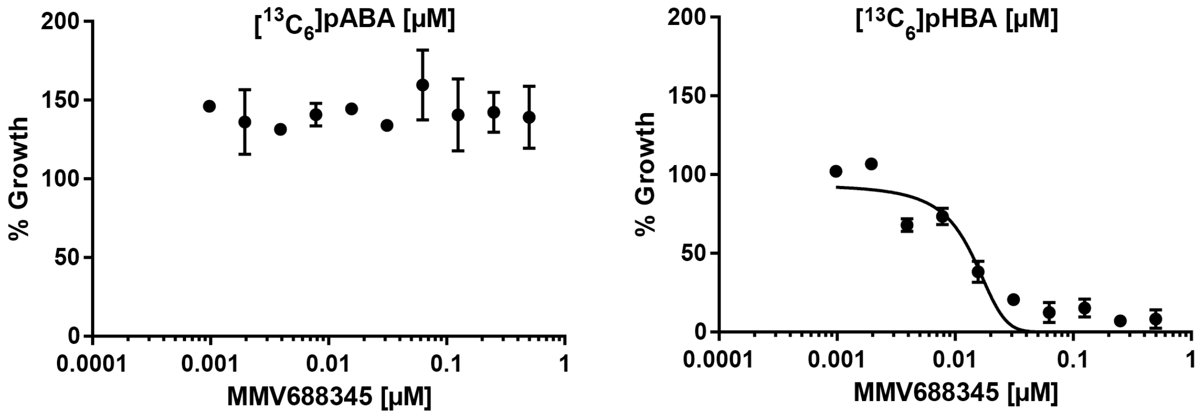


**Fig. S2. (continuation) Reversal of growth inhibition by MMV688345.** *P. falciparum in vitro* reversal of growth inhibition by MMV688345 in the presence of 7.3 µM [$^{13}C_6$]pHBA or [$^{13}C_6$]pABA was assessed in MM by SYBR green assay after 72 h incubation.

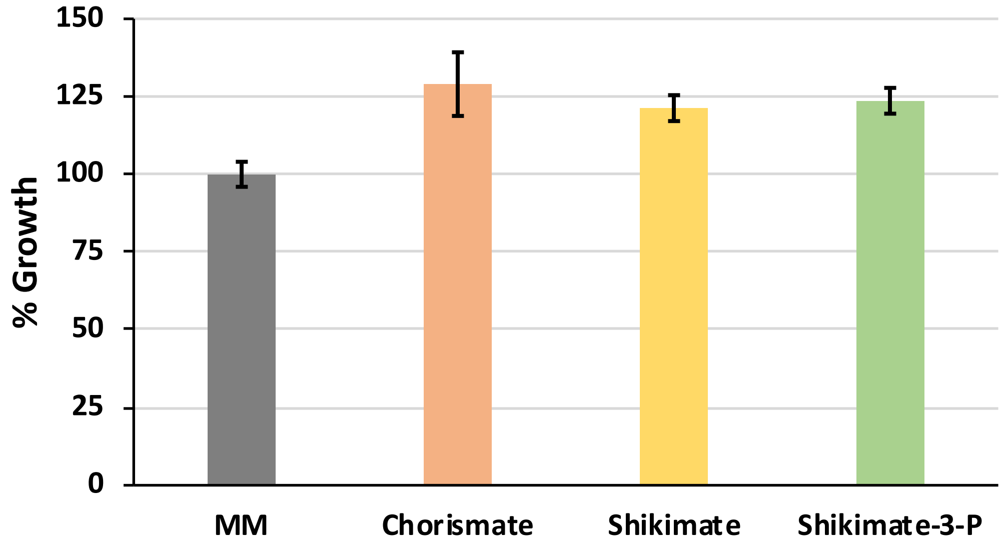


**Fig. S3. *P. falciparum* growth in MM in the presence of different metabolites from the shikimate pathway.** Cultures were supplemented with chorismate (12.5 µM), shikimate (25 µM) or shikimate-3-phosphate (25 µM) and growth was assessed by SYBR green assay. Values represent the mean ± S.E.M. from at least three independent assays performed in triplicate. *P* values for each supplemented metabolite with respect to control are: (*) < 0.00001, (**) < 0.000004 and (***) < 0.000009.

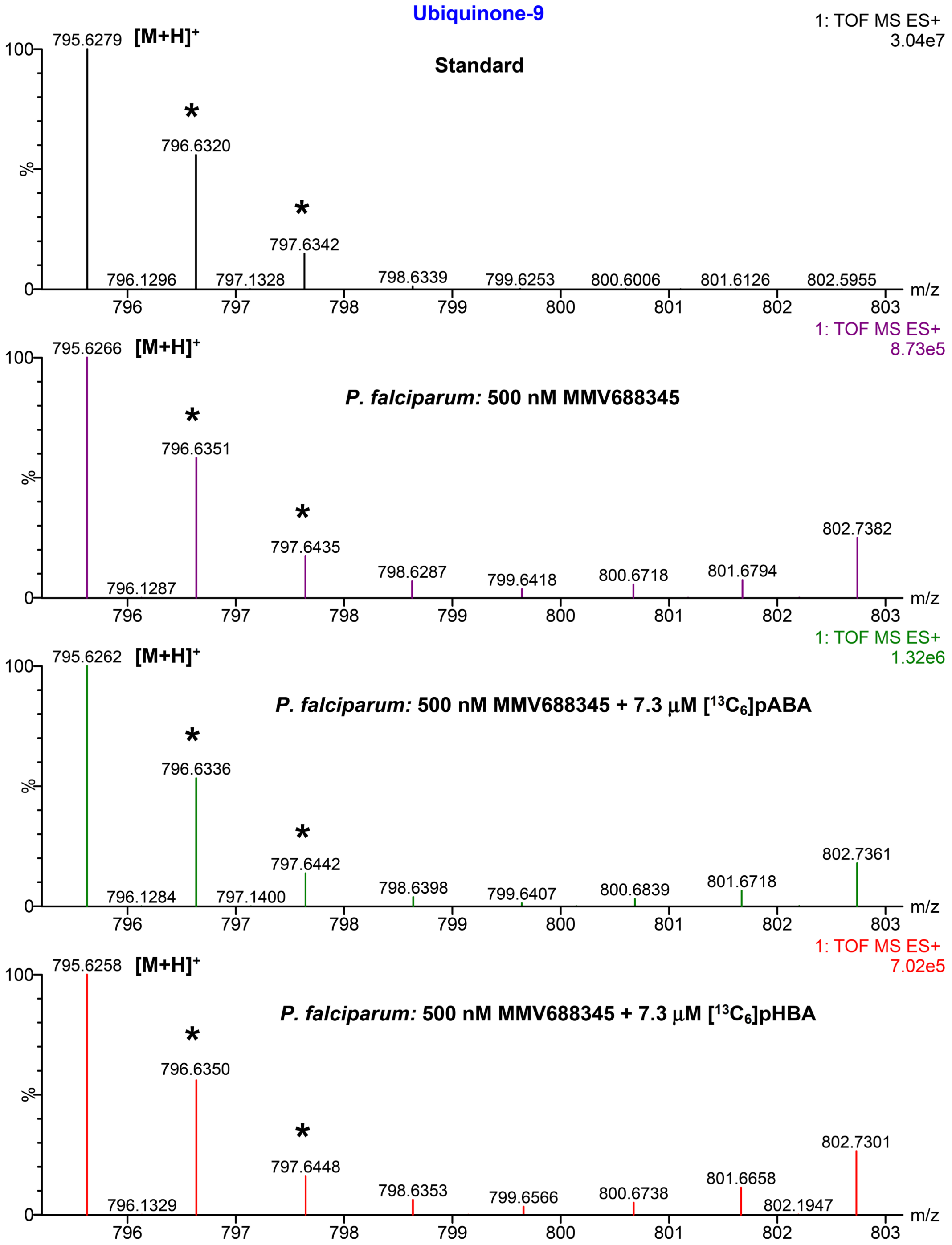


**Fig. S4. LC-HRMS positive-ion mode spectra of ubiquinone-9 to assess whether *P. falciparum* is able to use [$^{13}C_6$]pHBA and [$^{13}C_6$]pABA as a metabolic precursor for ubiquinone biosynthesis**. $[M+H]^+$ indicates the positive-ion corresponding to the mass of ubiquinone-9 and (*) indicates its natural isotopic distribution. The ion corresponding to the [$^{13}C_6$]ubiquinone-9 ($[M+H]^+$ expected = 801.6487) for [$^{13}C_6$]pHBA or [$^{13}C_6$]pABA incorporation into the head group of ubiquinone-9 was not detected.

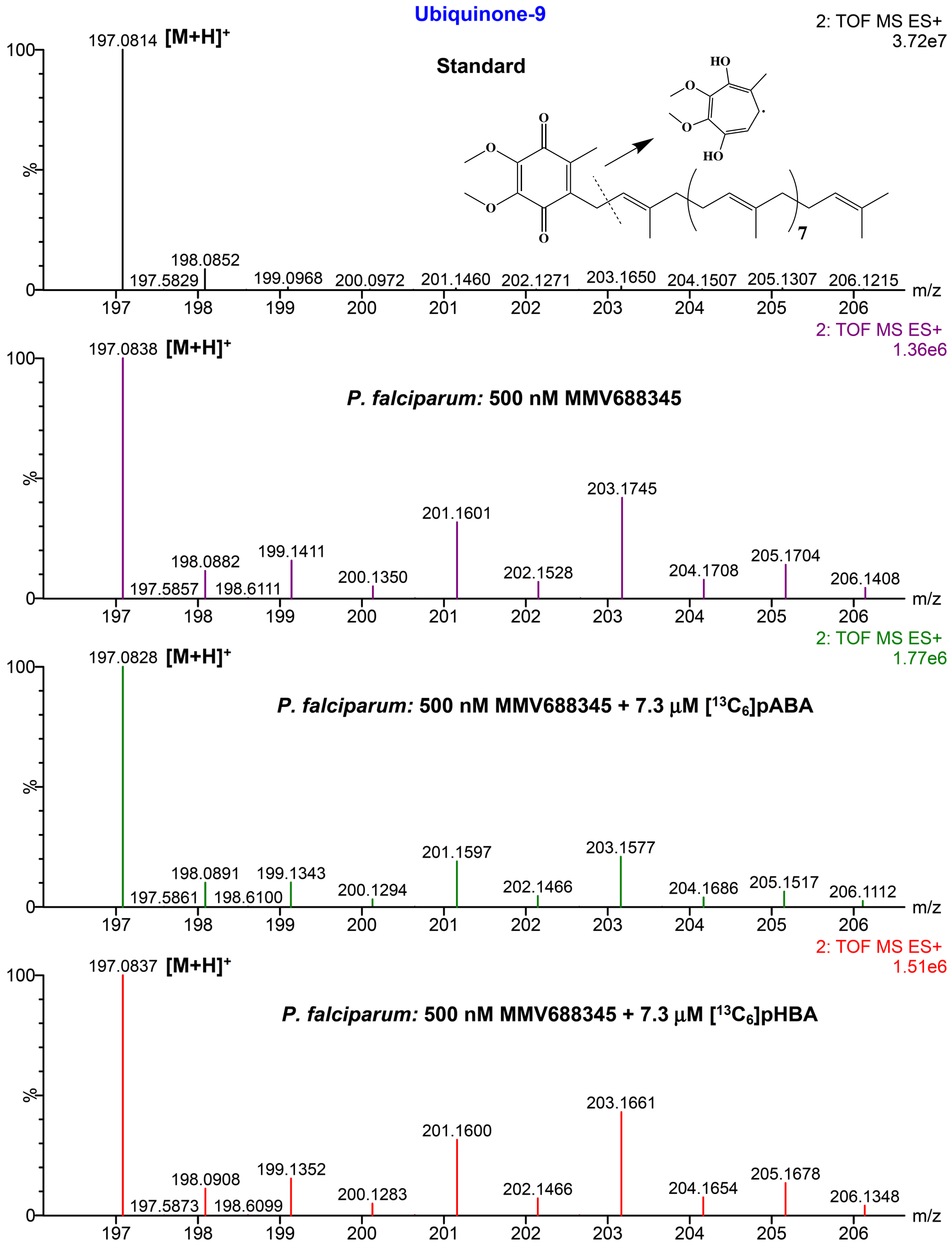


**Fig. S4. (continuation) Mass fragmentation profile of ubiquinone-9.** $[M+H]^+$ indicates the positive-ion corresponding to the mass of the ubiquinone-9 tropylium ion ($[M]^+$ expected = 197.0808). The predicted fragmentation is shown. The $[^{13}C_6]$tropylium ion ($^{13}C_6$-$[M]^+$ expected = 203.1010) was not detected.

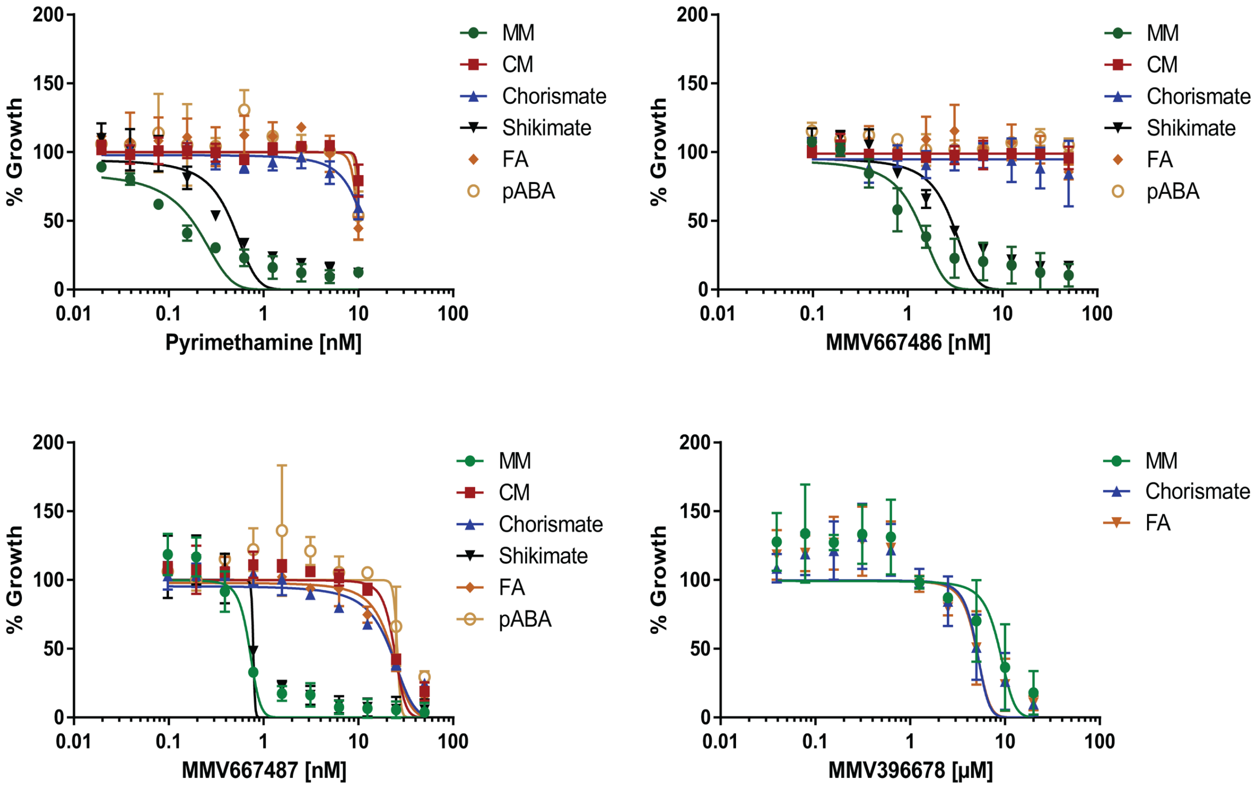


**Fig. S5.** Growth inhibition observed in MM by pyrimethamine ($IC_{50}$ = 0.18 ± 0.03 nM), MMV667486 ($IC_{50}$ = 1.28 ± 0.17 nM) and MMV667487 ($IC_{50}$ = 0.71 ± 0.05 nM) was reversed in CM while growth inhibition by MMV396678 ($IC_{50}$ = 8,723 ± 1.3 nM) was not reversed by FA. Chorismate, FA and pABA but not shikimate, also reversed growth inhibition by pyrimethamine, MMV667486 and MMV667487 similar to MMV688345. The following metabolite concentrations were used: 25 µM of shikimate, 12.5 µM of chorismate and 7.3 µM of FA.
